# Winners and losers over 35 years of dragonfly and damselfly distributional change in Germany

**DOI:** 10.1101/2020.08.03.234104

**Authors:** D.E Bowler, D. Eichenberg, K.J. Conze, F. Suhling, K. Baumann, A. Bönsel, T. Bittner, A. Drews, A. Günther, N.J.B. Isaac, F. Petzold, M. Seyring, T. Spengler, B. Trockur, C. Willigalla, H. Bruelheide, F. Jansen, A. Bonn

## Abstract

Recent studies suggest insect declines in parts of Europe; however, the generality of these trends across different taxa and regions remains unclear. Standardized data are not available to assess large-scale, long-term changes for most insect groups but opportunistic citizen science data is widespread for some taxa. We compiled over 1 million occurrence records of Odonata (dragonflies and damselflies) from different regional databases across Germany. We used occupancy-detection models to estimate annual distributional changes between 1980 and 2016 for each species. We related species attributes to changes in the species’ distributions and inferred possible drivers of change. Species showing increases were generally warm-adapted species and/or running water species while species showing decreases were cold-adapted species using standing water habitats such as bogs. We developed a novel approach using time-series clustering to identify groups of species with similar patterns of temporal change. Using this method, we defined five typical patterns of change for Odonata – each associated with a specific combination of species attributes. Overall, trends in Odonata provide mixed news – improved water quality, coupled with positive impacts of climate change, could explain the positive trend status of many species. At the same time, declining species point to conservation challenges associated with habitat loss and degradation. Our study demonstrates the great value of citizen science data for assessing large-scale distributional change and conservation decision-making.

## Introduction

Recent studies suggest long-term declines of insect populations in different parts of Europe (Conrad et al. 2006; Hallmann et al. 2017; Valtonen et al. 2017; Homburg et al. 2019). Such declines are a major conservation concern, especially because they could have a broad range of cascading effects for other species (Hallmann et al. 2014; Cardoso et al. 2020). However, many studies on insect change are based on local or small-scale datasets. Assessing change over large-spatial scales is difficult for most insect taxa due to a lack of standardized monitoring. Nonetheless, effective conservation policies strongly rely on knowledge of the large-scale trends of species. To facilitate conservation decision-making, there is an urgent need to make use of all available data to estimate large-scale and long-term changes in insect populations and communities.

While large-scale standardized insect monitoring is rare, at least beyond butterflies (van Swaay et al. 2008), large amounts of opportunistic data, without a common sampling protocol, are collected by citizen science (CS) (Chandler et al. 2017). CS data are associated with numerous statistical challenges but they have the advantage of a large spatial coverage and a reasonable time-span. Moreover, since some citizen scientists are active year-round, CS data also tend to capture a larger proportion of the biological community, including rare species, than standardized data (Bradter et al. 2018). As CS data has become more accessible, there has been simultaneous development of methods to analyze such data (van Strien et al. 2013; Isaac et al. 2014; Rapacciuolo et al. 2017). Occupancy-detection models have emerged as one approach that is robust to different ways citizen scientists collect data, by explicitly modelling heterogeneity in survey effort and species detectability (Isaac et al. 2014).

Odonata are a good case study for the application of occupancy-detection models to study long-term change since, despite a few exceptions, they are mostly not surveyed by structured monitoring over large spatial scales but are subject to extensive CS recording. Recent studies of dragonflies in different European countries suggest many species have increased (Powney et al. 2015; Termaat et al. 2019). In general, there is evidence of recent population increases of many freshwater organisms in Europe, thought to be due to better waste water treatment enabling recovery from previous impacts of water pollution (van Strien et al. 2016; Van Klink et al. 2020), especially following the implementation of the European Water Framework Directive (WFD) in the 2000s (Giger 2009; Hering et al. 2010). Climate changes signals have also been apparent by the northward expansion of southern species (Hickling et al. 2006; Ott 2010).

We used opportunistic data, including CS data, to study Odonata distributional change – both in terms of mean long-term trends and specific temporal patterns of change – since 1980 in Germany, which contains the largest number of Odonata species in Europe (Brockhaus et al. 2015; Kalkman et al. 2018). We further identified species attributes (i.e., species-specific characteristics) associated with distributional changes. Our analysis was part exploratory, to identify predictive attributes, and part hypothesis-driven, for some key attributes assumed to link to specific drivers. We hypothesized that sensitivity to climate change may be affected by species’ temperature preferences; while sensitivity to environmental management would be affected by species’ habitat preference. Specifically, we predicted increases of warm-adapted and early spring species and decreases of cold-adapted species, as well as increases of running water species, since these are thought to most benefit from improved water and environmental management (Termaat et al. 2015). We expected to see recovery of running water species especially during the 2000s when the WFD activities began. By contrast, increases associated with climate change were expected to be already visible from the 1980s. Finally, we tested whether communities had become more dominated by widespread and generalist species in line with biotic homogenization (Powney et al. 2015).

## Methods

### Data compilation

In collaboration with the Society of German speaking Odonatologists (GdO) and various conservation agencies representing different federal states, we compiled occurrence records on Odonata across Germany. The aggregated dataset comprised heterogeneous data, collected by both official and voluntary nature conservation organizations, without a common sampling protocol; however, experienced naturalists collected most of the data (Mauersberger et al. 2013; Trockur 2013; Petzold & Fritzlar 2014; Brockhaus et al. 2015). Available data usually included information on observer or project, date of observation, life stage of species (e.g. larva, exuviae, adult) and geographic coordinates or ordinance survey quadrant (Meßtischblatt Quadrant, MTBQ) of c. 5 × 5 km (Goertzen & Suhling 2019). Observations of adults comprised 93% of the data; hence, larvae data were excluded but exuviae were retained. In total, we compiled over 1 million records (Fig. 1).

**Figure 1:**
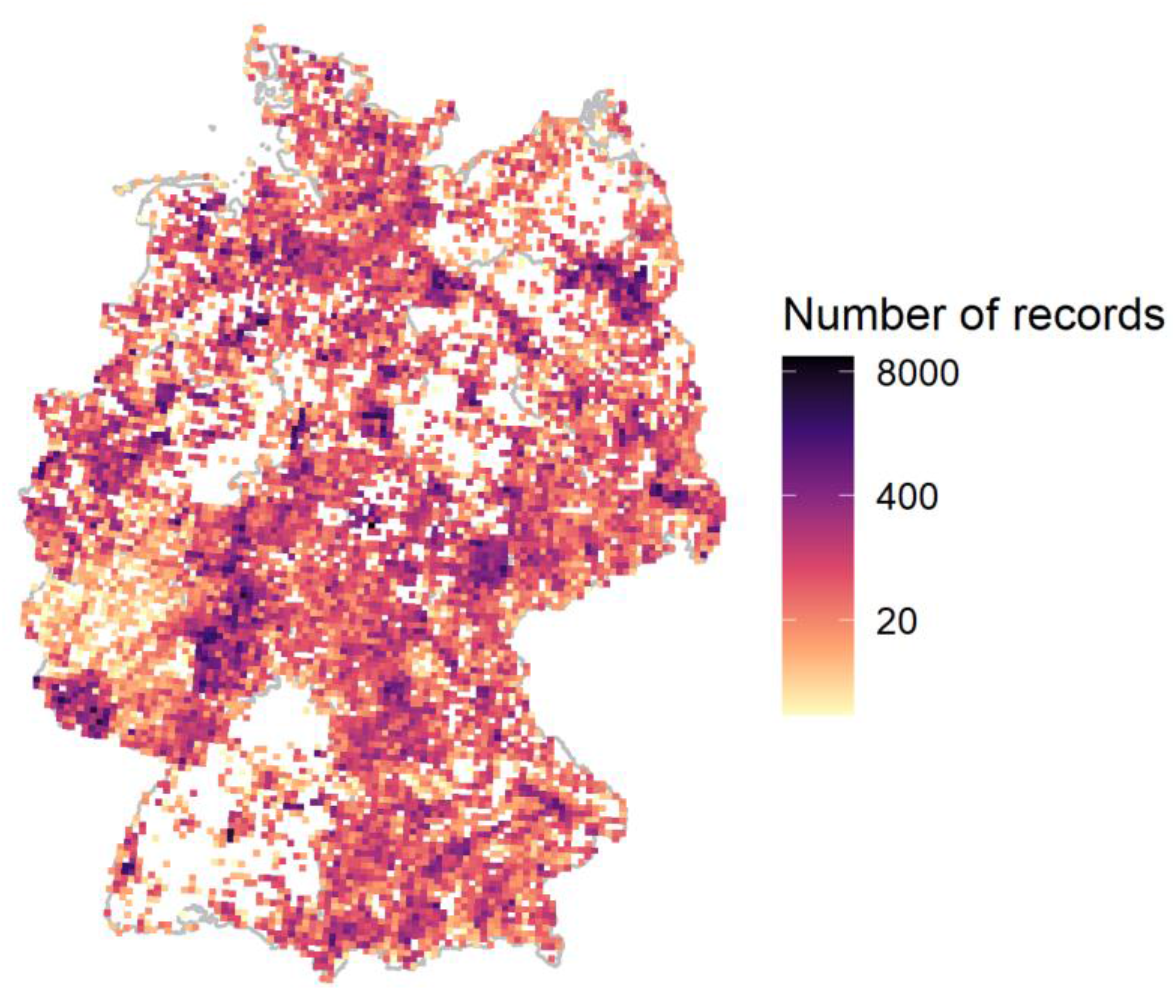
Distribution of Odonata occurrence records for each c. 5 × 5 km survey quadrant in Germany. Shading refers to the number of occurrence records. Data shown were subset to quadrats with records in at least 2 years since 1980.

### Data processing

We excluded: (1) data before 1980 since there were relatively few; (2) survey quadrant if they had not been visited in at least two separate years and (3) species seen in less than 25% of years (4 species: *Coenagrion hylas*, *Gomphus simillimus, Lestes macrostigma, Onychogomphus uncatus*) due to insufficient data to estimate a trend. We included seasonal migratory species, such as *Anax ephippiger* and *Sympetrum fonscolombii*, even though they do not overwinter in Germany. Overall, 81 species of Odonata were included in the last published atlas for Germany (Boudot & Kalkman 2015; Brockhaus et al. 2015) – our analysis included 77 species after applying the above exclusion criteria (Table S1). The finest unit of our analysis was a visit, referring to data collected on a given date in a given survey quadrant by a given observer/project. Therefore, we organized the data into a list of species records seen on a given visit. As a measure of sampling effort, we calculated the total number of recorded species (often called “list length”) on each visit.

### Attribute data

We selected attributes that reflect species’ exposure, sensitivity or adaptive capacity to environmental changes, such as climate change, land-use or conservation measures.

#### Distribution

We estimated species’ European geographic range size as the number of occupied grids in a recent atlas (Boudot & Kalkman 2015).

#### Species temperature preference

Species’ temperature preferences were calculated by overlaying each species’ European distribution with an average temperature map from E-OBS v. 19e (Cornes et al. 2018) following other studies (Jiguet et al. 2007). For each species, we calculated the mean of the mean daily temperatures of occupied grid cells. While we call this variable “temperature preference”, its calculation did not aim to estimate species’ optimal temperatures but rather to place species on a gradient from those preferring cooler temperature to those preferring warmer temperatures.

#### Life-history

Data on voltinism, i.e., number of generations per year, was compiled from Corbet et al. (2006). We applied a weighted mean of fuzzed-coded species affinities (summed to 10) to different categories: multivoltine (coded as 5), bivoltinie (4), univoltine (3), semivoltine (2), and partivoltine (1). This weighted mean ranged between 1 (for a fully partivoltine species) and 5 (for a fully multivoltine species).

#### Phenology

Mean start dates of the main flight period were taken from Boudot & Kalkman (2015). Species’ phenologies vary geographically but the data presented was usually for Bavaria, southern Germany. However, like for temperature preference, the aim was to only create a variable that placed species on a gradient from those appearing early in the year to those appearing later.

#### Habitat

Main habitat preferences were classified according to descriptions in Dijkstra (2006) and Boudot & Kalkman (2015). Each species was coded to whether they use the following habitats: streams, rivers, ponds, lakes, ditches, canals, fenland, bogs, forest and quarries/pits.

#### Morphology

Wing length (median of lower and upper values) was taken from Dijkstra (2006).

#### Threat-level

We compiled data on the 2015 red list classification for each species in Germany (Ott et al. 2015). We aligned the German categories with the international IUCN categories following Jansen et al. (2020).

Attribute data are provided in Data S1.

### Annual occupancy estimates

We used occupancy-detection models that account for imperfect detection, which have been used in previous studies using opportunistic CS data (van Strien et al. 2013; Outhwaite et al. 2020) and tested in simulation studies (Isaac et al. 2014). Imperfect detection occurs when a species inhabits a site but is not detected by the observer during a visit. Detection probability here also includes recording probability (i.e. the citizen scientist does not necessarily record all species that they detect). Detection probability of species is estimated by making an assumption about closure: a period during which species’ occupancy does not change. The number of times a species was/was not seen during this period of closure informs on detectability. We estimated the flight season of each species as between the lower 5% and upper 5% of days of year when each species was seen across all records. We only fit the model for each species to the subset of records during the flight season to meet the closure assumption. Models were fit to each species separately.

Letting *Z*_*i,t*_ refer to the true occupancy status for a species in a survey quadrat *i* in a year *t*, we modelled occurrence probability (*ψ*) as a function of site and year variation. Year variation was modelled by including year as a fixed effect factor. Site variation was modelled as a series of random terms: ecoregion variation (‘Naturräumliche Großregionen Deutschlands’) (Bundesamt für Naturschutz 2008) at two spatial scales (level 1: 7 coarser-scale regions and level 2: finer-scale 487 regions) and survey quadrant variation:

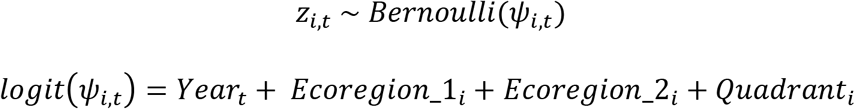

Detection probability (*p*) was modelled for each visit *j* to a given quadrant in a given year, and assumed to depend on year, day of year (accounting for species’ phenology) and survey effort (a single list with 1 species record, a short list with 2–3 species records, or a longer list).

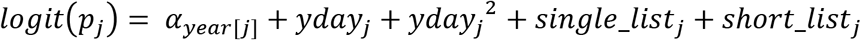

The observed detection data for a given species, *y* (0 or 1), on each visit are then assumed to be drawn from a Bernoulli distribution conditional on the presence of the species in that quadrant and year:

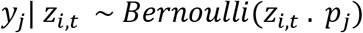

We also ran dynamic-occupancy models that explicitly model changes in occupancy between years as either caused by persistence probability when the site was occupied in the previous year or by colonization probability when the site was not occupied in the previous year. For most species, these models produced similar patterns to the simpler occupancy model above; but for the rarer species, the parameters of the dynamic-occupancy model showed lower convergence and large uncertainty, especially for the earlier years when there was less data. Hence, we used our simpler occupancy model that produced satisfactory results for the larger number of species.

Using the predicted occurrences (*Z*_*i,t*_), we calculated a number of derived parameters: (1) the proportion of survey quadrants that were occupied by a species in each year (hereafter ‘occupancy proportion’); (2) the slope of a regression line through the annual occupancy proportions for each species (hereafter ‘trend’) and (3) proportional change in occupancy proportion between the first and last years. These statistics were calculated during model fitting and hence the uncertainty in occupancy predictions was retained in each parameter.

The models were fit by Bayesian inference using JAGS, a program for fitting hierarchical models using Markov Chain Monte Carlo simulation. We used vague priors for most parameters but a random walk prior, to share information across years, for the year effect on occupancy (Outhwaite et al. 2018). We used 3 chains with 50000 iterations, discarding the first 25000 as burn-in. Model convergence was assessed by the Rhat statistic and traceplots. The model code is provided in the Supplementary Material.

### Long-term trends

We analyzed the inter-specific variation in distribution trends using a linear regression model, with species attributes as explanatory variables and species’ trend estimates as the response. Correlations among species’ attributes were examined to check for any possible collinearity issues. We first conducted an exploratory analysis with each of the habitat variables in simple regression models. Using these models, we identified habitat variables that explained variation in trends (i.e., if p<0.05). We then combined the selected habitat variables along with the other variables (temperature preference, voltinism, flight start date and wing length) in a multiple regression model of long-term trends. We performed step-wise deletion, removing insignificant attributes, to identify the best model. We also used a linear model to compare the trends of species with different red list status.

Since species do not necessarily provide independent data points due to shared evolutionary histories, we checked whether the results of the regression were consistent after accounting for species relatedness. We used the taxonomic classification to build a simple phylogenetic tree with equal branch lengths for each taxonomic rank. We then included the tree as a correlation structure (corPagel ‒based on Brownian motion) in a generalized least squares model (Paradis & Schliep 2019). Since this had little effect on the effect sizes of the attributes, and the estimated phylogenetic signal was close to zero (likelihood ratio test between models with and without the correlation structure, p=0.15), we present the simpler model without this correlation structure.

### Temporal patterns

We used a time-series clustering method to group together species showing similar dynamics using the TSclust package (Montero & Vilar 2014). The mean estimates of annual occupancy proportions were preprocessed via logit transformation. We used the Pearson correlation coefficient as a dissimilarity measure and clusters were split using partitional clustering. We selected the number of clusters using several cluster indices (average silhouette width, dunn index, separation index), assessed over the range of 2 to 20 clusters. We visualized the temporal change in each pattern by calculating the geometric mean of species’ occupancy proportions in each cluster. To include the uncertainty of species’ occupancies, we also combined 1000 random draws from the posterior of the annual occupancy estimates for the species in each cluster and calculated the mean and upper and lower 2.5% quantiles.

### Assemblage-level properties

To examine the implications of the species-level changes for the total Odonata species pool (referred to here as assemblage-level), we aggregated the predicted occupancy proportions of all species. We calculated for each year: mean species richness and diversity (Shannon index) based on species’ occupancy proportions and community-weighted means for range size (mean range size weighted by occupancy proportion). We retained the uncertainty of the species’ estimates by repeating the calculations for 1000 random draws from the posterior distributions of the species’ occupancies and calculating the mean and upper and lower 2.5% quantiles.

We used R version 3.6.3 for all analysis. Statistical significance was assessed when 95% confidence intervals did not overlap zero.

## Results

### Long-term trends

Of the 77 species, we found that 37 species were significantly increasing and 18 were significantly decreasing (Fig. 2A). Of the species with no significant trend, 20 species had a small trend estimate (within 5% long-term change) while 2 species had a larger mean trend estimate. Species with both a large positive trend and undergoing large range expansions included: *Crocothemis erythraea* (increased in occupancy by a factor of 84, from 0.3% to 27% of occupied survey quadrants) and *Erythromma viridulum* (increased by a factor of 8) (Fig. 2B). By contrast, species with large negative trends and undergoing large range contractions included: *Sympetrum danae* (decreased by a factor of 0.40, from 79% to 32% of occupied survey quadrants) and *Coenagrion hastulatum* (decreased by a factor of 0.28) (Fig. 2B). The proportions of dragonflies versus damselflies increasing (25/50 vs 12/27) and decreasing (9/50 vs 9/27) were not significantly different (chisq test, p=0.4). Estimated population trends were consistent with the 2015 red list classification. Threatened species (vulnerable, endangered, critically endangered) had less positive population trends than species of least concern or near threatened (P<0.05, Fig. 2C).

**Fig. 2:**
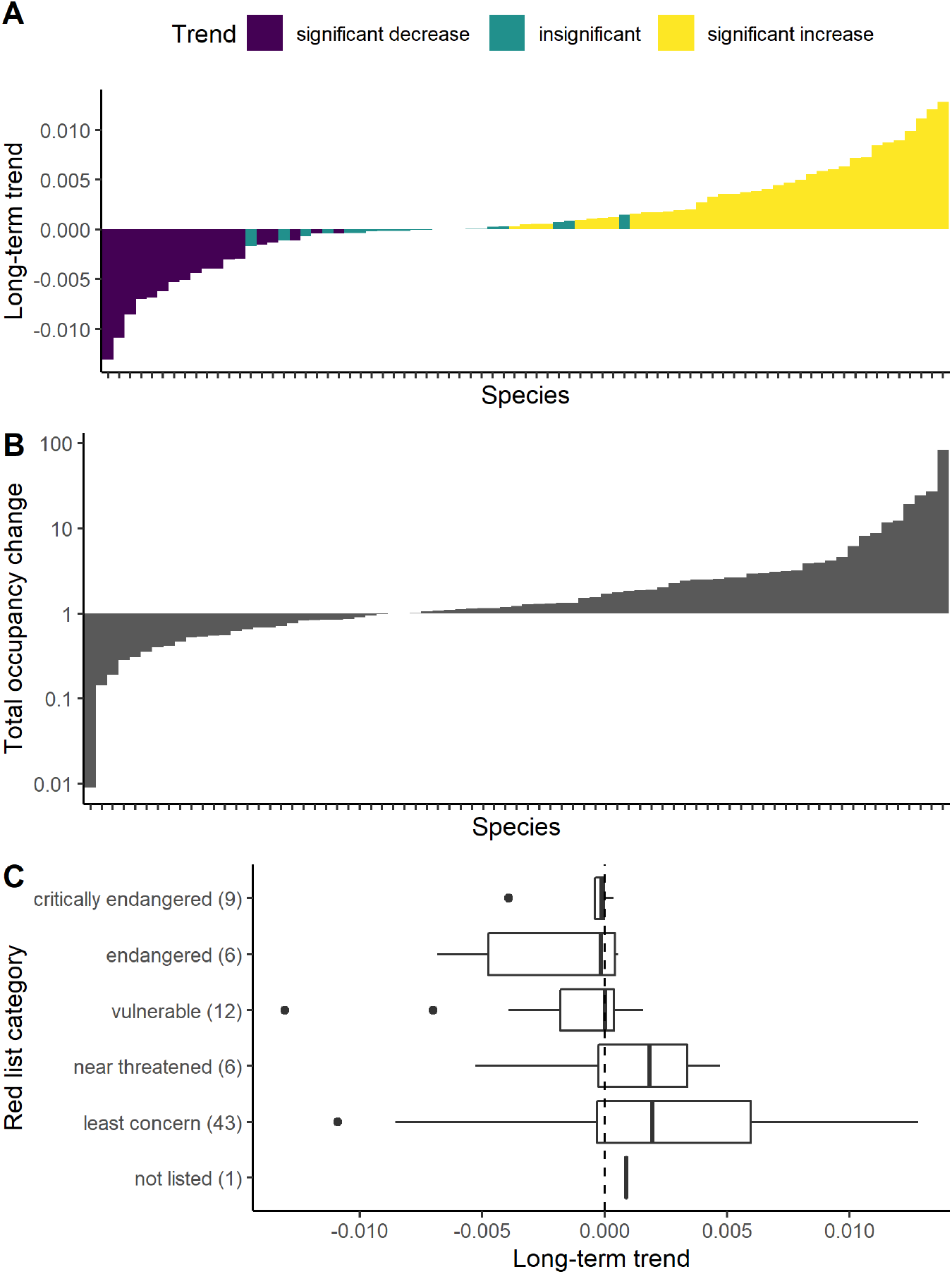
Estimated nationwide trends in Odonata species’ distributions. (A) species’ long-term trends (0 = no trend), ordered by their magnitude of trend; (B) ratio in the estimated number of occupied grids between the last (2016) and first (1980) year (ordered by their magnitude of change, 1 = no change) and (C) boxplots (outliers are showed as solid points) of the association between long-term trend and red list classification (number of species in each group is shown in brackets). See Fig. S1 for time-series of individual species and Data S1 for data on the trends of each species.

In the analysis of habitat associations, species using river habitats tended to increase; while species associated with bog habitats tended to decrease (Fig. 3). In a multiple regression model, the most important attributes explaining variation in species’ long-term trends were temperature preference, flight start date, wing length and river use (Fig. 4). Temperature preference was positively associated with species’ trends: warm-adapted species increased while cool-adapted species decreased (Fig. 3). Species that appear as flying adults earlier in the year also had more positive population trends. Wing length and river habitat use were also positively associated with trends (Fig. 3). While bog use was negatively associated with species’ trends in the simple regression model, it was no longer significant in the multiple regression model (Fig. 4). This was probably because bog species were associated with colder temperature preferences (r = − 0.54), leading to a more uncertain effect of bog use after accounting for temperature preference (Fig. 4). Voltinism had little importance (Figs 3 & 4). Together, temperature preference, wing length, flight start date and river use explained 29% of the variation in trends among species.

**Fig. 3.**
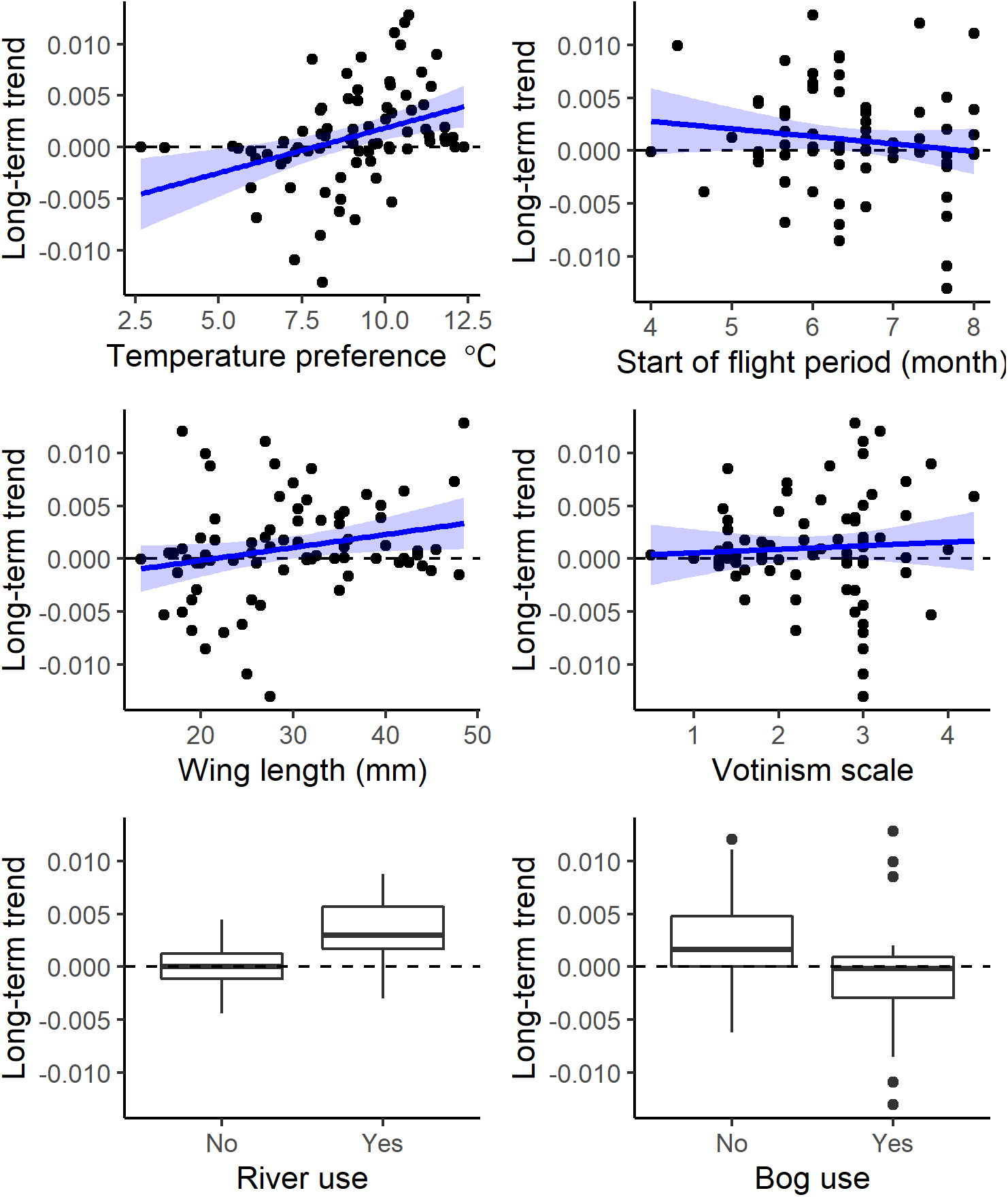
Relationships between each attribute and species’ population trends, each point is a species. The blue line is a fitted simple regression line. Boxplots are shown for river and bog use. The dashed horizontal line is the line of stable trends.

**Fig. 4:**
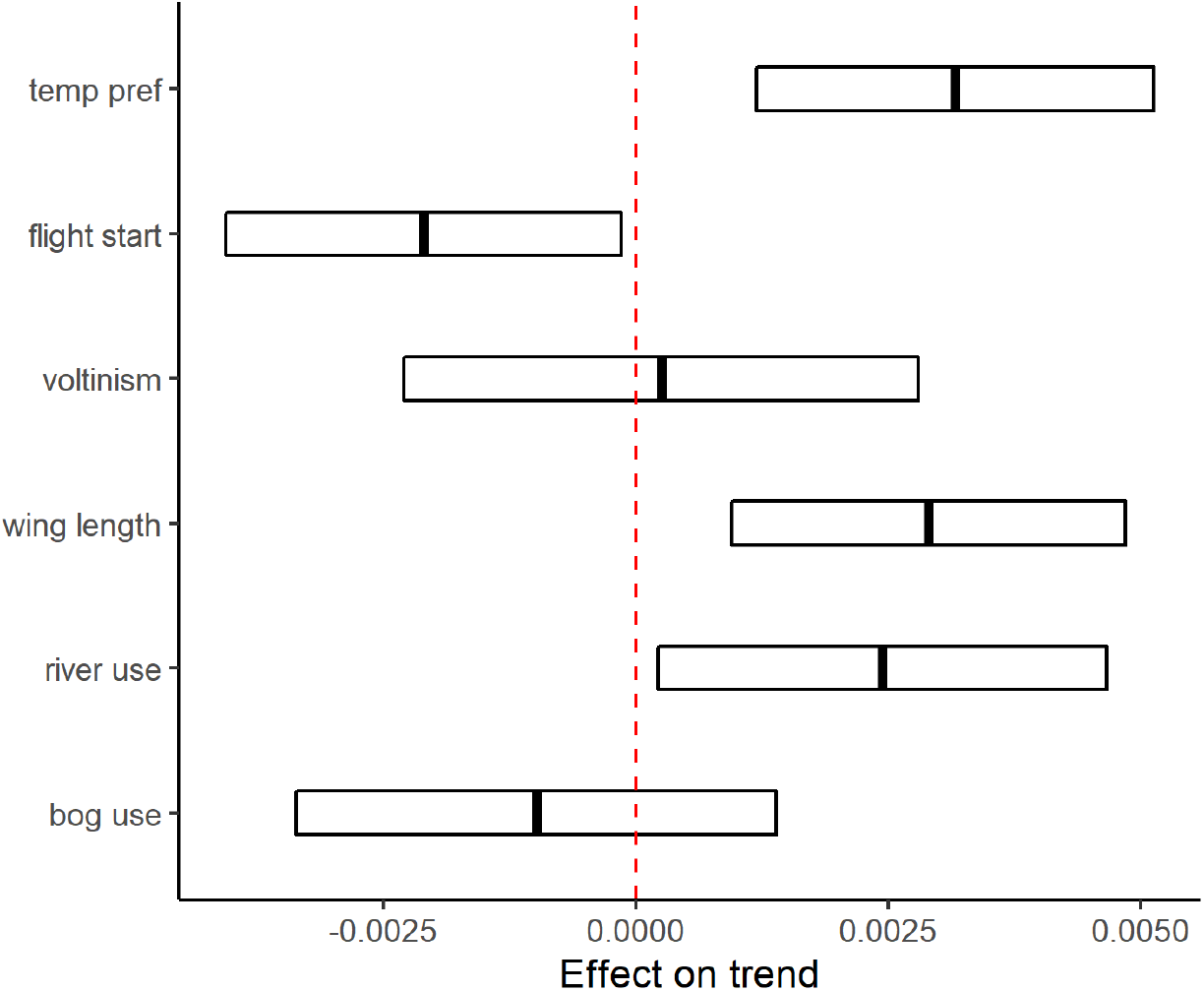
Effect of species’ attributes on their long-term distribution trends tested in a multiple regression model. Continuous variables (all except river and bog use) were scaled to units of 2 standard deviations to facilitate comparison with the binary habitat variables. The dashed red line is the line of no effect. Shown are the mean effect ± 95% confidence interval.

### Analysis of temporal patterns

Time-series clustering of species’ occupancy dynamics resulted in five grouping of species with similar patterns of change (Fig. 5). Despite the different temporal patterns, clusters 1 and 2 and clusters 3 and 4 did not have significantly different long-term trends. Cluster 1, the largest group of 34 species, comprised species with a persistent increase in occupancy but with a decline in the last years (Fig. 5A). Warm-adapted riverine species were most likely to be within this first group (Fig. 5B). Cluster 2 included 11 species that were initially stable but increased since the 2000s – this group included some of the most warm-adapted species (Fig. 5). Cluster 3 included 5 species, all using bog habitats, which showed variable trends but were typically declining mid 2000s onwards. Cluster 4 included 7 species that declined during the 1980s and 90s and tended to be cold-adapted bog species (Fig. 5). Finally, cluster 5 comprises 20 species with a mostly consistent decline and tended to be cold-adapted and short-winged species (Fig. 5).

**Fig. 5:**
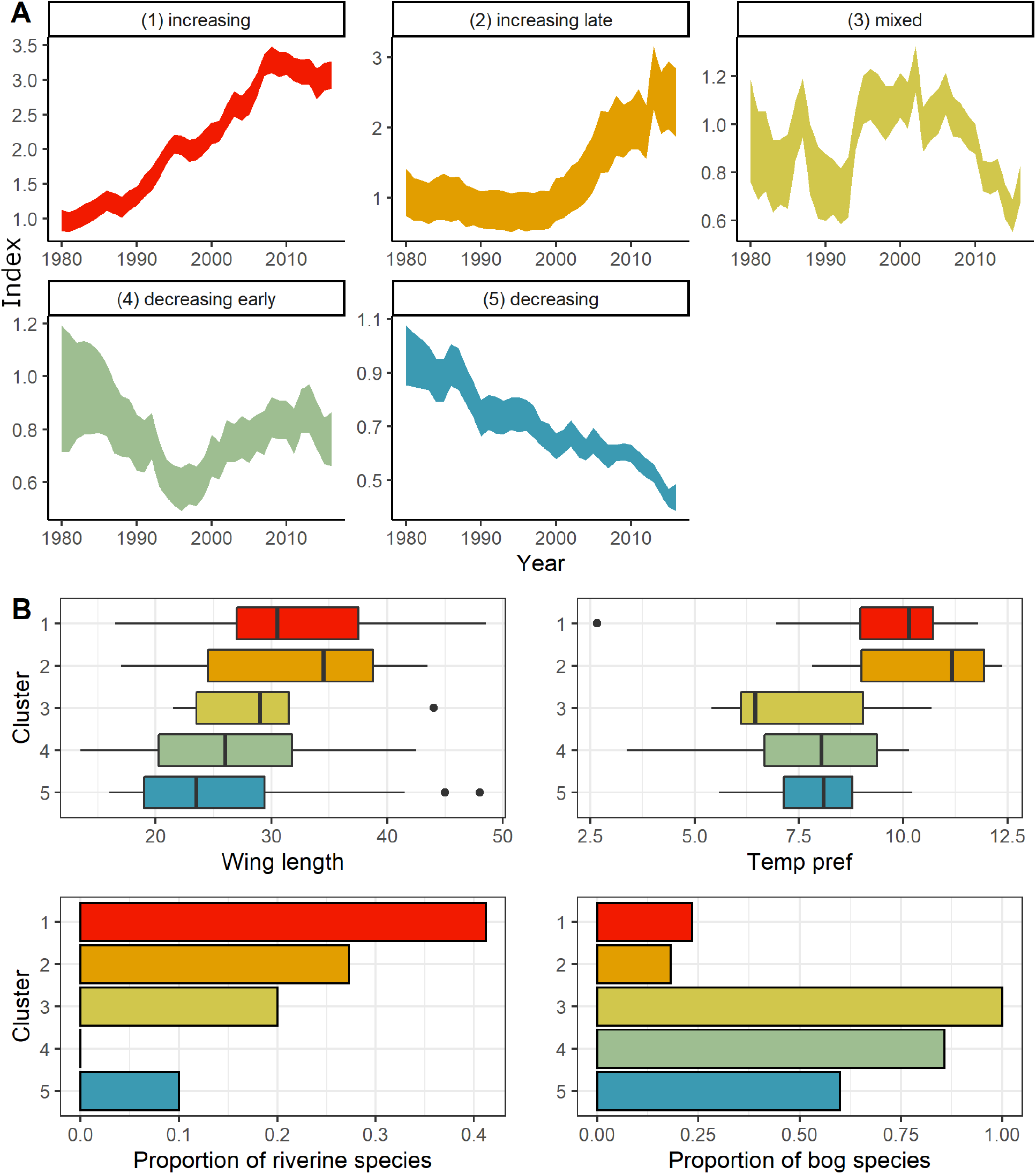
Time-series clusters and associated attributes. (A) Each cluster reflects a common pattern of change in occupancy over time for a group of species. The index represents the annual mean occupancy estimate relative to 1980. The number of species in each cluster were: 34, 11, 5, 7 and 20. (B) The plots below shown boxplots or barcharts for attribute values within each cluster. See Table S2 for list of species in each cluster. (A) was repeated by removing species with the most extreme occupancy changes in each cluster to check they were not driving the patterns the mean patterns – but similar patterns were found (Fig. S2).

### Assemblage-level consequences

Predicted mean species richness per quadrat has generally increased over the time period; however, some periods of decline were apparent, especially during the 1980s and during the 2010s (Fig. 6). Diversity shows similar trends, with increases until 2010 and decreases since then. Time-series of the weighted mean in range size suggested no simple shift towards widespread species – larger-range species became more dominant in the 1980s but smaller-range species became more dominant in the 1990s and 2000s.

**Fig. 6:**
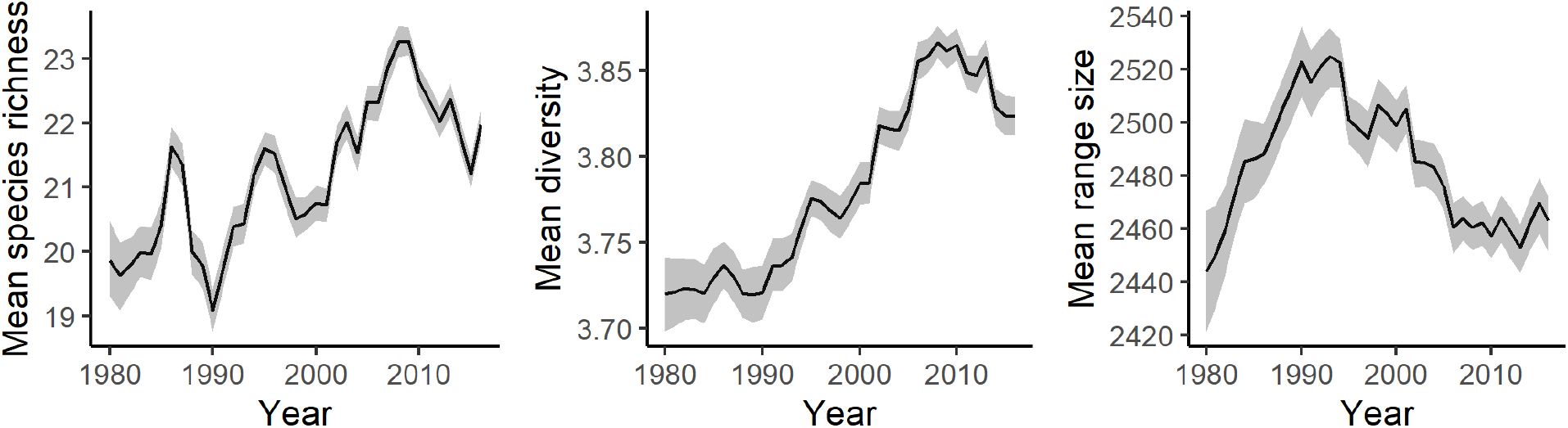
Time-series of aggregate predictions across all species: mean species richness and diversity per survey quadrat, and weighted mean of European range size. Shown are means and 95% CI of the mean.

## Discussion

Freshwater habitats have faced multiple anthropogenic threats, including eutrophication, acidification, climate change and canalization (Vörösmarty et al. 2010). Globally, freshwater vertebrate species are reported to be declining (He et al. 2019). Hence, our findings of many stable or increasing Odonata species since 1980 in Germany might seem surprising. However, our results are consistent with other studies on Odonata species that show increasing trends in Europe (Powney et al. 2015; van Strien et al. 2016; Termaat et al. 2019) and more generally positive trends in biomass of freshwater insects (Van Klink et al. 2020).

Climate change probably plays a key role in the success of many Odonata species in Europe. As highly-mobile organisms, many Odonata species may have adaptive capacity to respond to climate change, demonstrated by range-shifts reported in other countries (Hickling et al. 2005; Braune et al. 2008; Flenner & Sahlen 2008), and may be responding stronger to climate change than many other terrestrial species (Hassall 2015). We find that formerly rare warm-adapted species such as the scarlet darter, *Crocothemis erythraea*, and the small red-eye damselfly, *Erythromma viridulum*, have undergone large range expansions across Germany. Increases of species typically appearing earlier in the year might also be linked to warming temperature. Earlier and longer reproduction seasons may increase the potential for more than one generation within a year (Braune et al. 2008) or allow early breeders to monopolize resources ahead of later-breeding species (Suhling & Suhling 2013). Additionally, we found that wing-length was an important predictor. One of the biggest winners has been the Emperor, *Anax imperator*, a strong flier with long wings (Ruppell 1989).

Our findings may also reveal the impacts of improved environmental management, especially for rivers. Many running water species were increasing 1980 onwards even though the EU Water Framework Directive (WFD), which aimed to improve water quality, was not adopted until 2000 (Hering et al. 2010). This is probably because there were a range of other conservation and environmental management projects to improve water quality prior to 2000 in Germany (Detering 2000; Giger 2009). These projects included expansion of water purification plants; improved watercourse management (less removal of vegetation and disturbance of sediments); Fauna-Flora-Habitat Directive activities that targeted specific species and conservation measures to improve degraded wetlands. Positive trends of dragonflies in the Netherlands were also thought to partly reflect habitat improvements (Termaat et al. 2015). River restoration projects in Europe have also allowed some recovery of other taxa, such as fish, though not necessarily to former historic states (Thomas et al. 2015). The success of riverine species may also reflect some synergism in the impacts of climate change and environmental management, since improvements in water quality may have facilitated climate change-driven range-expansion by increasing the establishment success of immigrants (Braune et al. 2008).

Despite improvements in some freshwater habitats, we also identified a significant number of declining, ‘loser’ species. Decreases of cold-adapted species could represent range contraction due to physiological stress under warming climates; however, more likely, they are a consequence of habitat loss, associated with climate change and land-use. Some declining species, such as the black darter, *Sympetrum danae*, and the northern damselfly, *Coenagrion hastulatum,* are cold-adapted and typical of bog or moorland habitats, which are among the most threatened habitats in Europe (European Environment Agency 2015). Overall, species of standing water habitats had more negative trends than those of running water habitats. While some types of standing water habitats, such as gravel quarries/sand pits, might have become more common, especially in eastern Germany, decreases in groundwater level due to overexploitation of water resources has probably reduced the availability of many small standing waterbodies. Small or shallow waterbodies have been also vulnerable to droughts (Opitz et al. 2019). The success of on-going conservation projects to restore bogs and other standing water habitats requires on-going monitoring (Dolny et al. 2018; Krieger et al. 2019).

Many studies on biodiversity change have focused on the simple long-term mean trend of a species but this approach can mask a diversity of more complex temporal responses (Outhwaite et al. 2020). Indeed, recent analyses of invertebrate changes in the UK found both time-periods of increases and time-periods of decreases, which varied among taxa (Macgregor et al. 2019; Outhwaite et al. 2020). For instance, UK freshwater organisms decreased between 1970 and the mid-1990s but increased from then until 2010 (Outhwaite et al. 2020). Moreover, simple trend models can produce trend estimates driven by fluctuations in particular years (Seibold et al. 2019). Using time-series clustering, we defined five characteristic patterns of change that were common across multiple species and which separated species that differed in paths of change, even when their long-term mean trend was similar. While there are other approaches to visualize the time-series patterns for specific groups of species, often called multi-species indicators (e.g., the Farmland Bird Index), species are pre-assigned to groups with current methods and the resulting indicators can be sensitive to grouping decisions (Gregory et al. 2019). Here, we show how it is possible to allow species to naturally group based on the similarity of their dynamics, which has the potential to reveal the role of previously overlooked community changes and may be less prone to subjective species selection decisions. Our approach may be used to develop an alternative sets of multi-species indicators that represent the multi-facetted nature of change within communities.

Despite evidence of species turnover, we found no trend towards biotic homogenization nor overall loss of diversity. Nonetheless, our findings have various implications for freshwater conservation in Germany. The strong associations between species attributes and population trends support the use of trait-based vulnerability assessments for conservation decision-making (Conti et al. 2014). However, increasing populations for some species represent expansion into new regions; while for other species, increases rather reflect recovery to formerly occupied parts of the range. The former group may be regarded as ‘neonatives’ (Essl et al. 2019) and contribute to the development of novel species interactions and assemblages (Carrasco et al. 2018), with currently unclear repercussions for established/native species (Flenner & Sahlen 2008; Suhling & Suhling 2013). Decreasing populations of other species have led to a decline mean in species richness during the 2010s. Standstills in the recovery of Odonata have been reported in the Netherlands (van Grunsven et al. 2020) and in the UK (Outhwaite et al. 2020). On-going monitoring and synthesis of habitat assessments is needed to assess the likely cause.

Since our analysis is based on opportunistic data, concerns should always remain about the robustness of the trends. The occupancy models aimed to account for observational processes but we cannot rule out changes in survey efforts affecting some of our results. Also, our analysis only focused on changes in distributions and not changes in abundance. For some species, increases or decreases in abundance may not yet have translated into changes in occupancy. We also only examined changes from 1980 onwards – most likely more historical data would highlight the earlier negative impacts of past water pollution and enable assessment of how much species have been able to recover from former historical ranges (Goertzen & Suhling 2019; Outhwaite et al. 2020).

Using an extensive citizen science dataset, our analysis revealed a complex picture of positive and negative, linear and non-linear distribution changes of Odonata species in Germany over the past 35 years. Climate change, habitat change and environmental management have probably all played a role. Cold-adapted habitat specialists of standing water habitats are likely to be most vulnerable to further environmental change; while increases of species associated with river habitats signal the conservation successes that can be achieved by better environmental management. Overall, our study demonstrates the value of the intensive recording efforts of citizen scientists in the past and highlights the need to support these efforts in the future, especially given signs of on-going Odonata declines over the last decade.

## Acknowledgements

This analysis was made possible through the efforts of many volunteer recorders over the past decades to whom we are very grateful – especially from the GdO - Gesellschaft deutschsprachiger Odonatologen e.V., but also from all the working groups at the level of the German federal states. We thank the various Arbeitskreis Libellen across Germany, and Michael Frank, Rüdiger Mauersberger, Frank Fritzlar and Samuel Rauhut, and the Thüringer Landesamt für Umwelt, Bergbau und Naturschutz for also sharing Odonata data for our project. We much appreciate the support of the German Research Foundation (DFG) for funding the sMon working group (Trend analysis of biodiversity data in Germany) through the iDiv (DFG FZT 118) synthesis center sDiv.

