## Supplementary Material for "Winners and losers over 35 years of dragonfly and damselfly distributional change in Germany"

Table S1: Species included in analysis

```
 Aeshna affinis               Aeshna caerulea
 Aeshna cyanea                Aeshna grandis
 Aeshna isoceles              Aeshna juncea
 Aeshna mixta                 Aeshna subarctica
 Aeshna viridis               Anax ephippiger
 Anax imperator               Anax parthenope
 Boyeria irene                Brachytron pratense
 Calopteryx splendens         Calopteryx virgo
 Ceriagrion tenellum          Chalcolestes viridis
 Coenagrion armatum           Coenagrion hastulatum
 Coenagrion lunulatum         Coenagrion mercuriale
 Coenagrion ornatum           Coenagrion puella
 Coenagrion pulchellum        Coenagrion scitulum
 Cordulegaster bidentata      Cordulegaster boltonii
 Cordulia aenea               Crocothemis erythraea
 Enallagma cyathigerum        Epitheca bimaculata
 Erythromma lindenii          Erythromma najas
 Erythromma viridulum         Gomphus flavipes
 Gomphus pulchellus           Gomphus vulgatissimus
 Ischnura elegans             Ischnura pumilio
 Lestes barbarus              Lestes dryas
 Lestes sponsa                Lestes virens
 Leucorrhinia albifrons       Leucorrhinia caudalis
 Leucorrhinia dubia           Leucorrhinia pectoralis
 Leucorrhinia rubicunda       Libellula depressa
 Libellula fulva              Libellula quadrimaculata
 Nehalennia speciosa          Onychogomphus forcipatus
 Ophiogomphus cecilia         Orthetrum albistylum
 Orthetrum brunneum           Orthetrum cancellatum
 Orthetrum coerulescens       Oxygastra curtisii
 Platycnemis pennipes         Pyrrhosoma nymphula
 Somatochlora alpestris       Somatochlora arctica
 Somatochlora flavomaculata   Somatochlora metallica
 Sympecma fusca               Sympecma paedisca
 Sympetrum danae              Sympetrum depressiusculum
 Sympetrum flaveolum          Sympetrum fonscolombii
 Sympetrum meridionale        Sympetrum pedemontanum
 Sympetrum sanguineum         Sympetrum striolatum
 Sympetrum vulgatum
```

Figure S1: Time series for all species (on same y-axis – see next page for same graph in which this is relaxed)

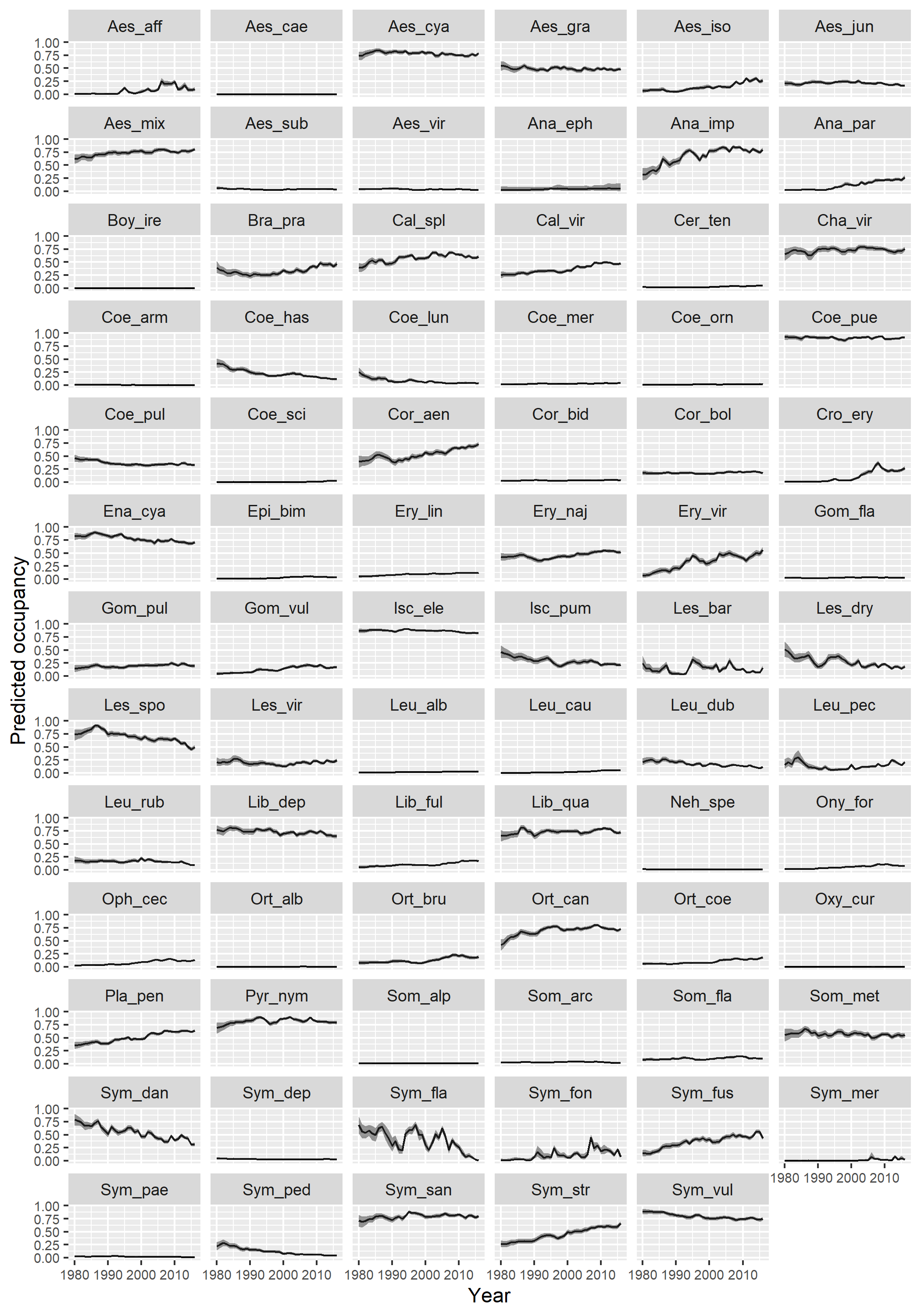

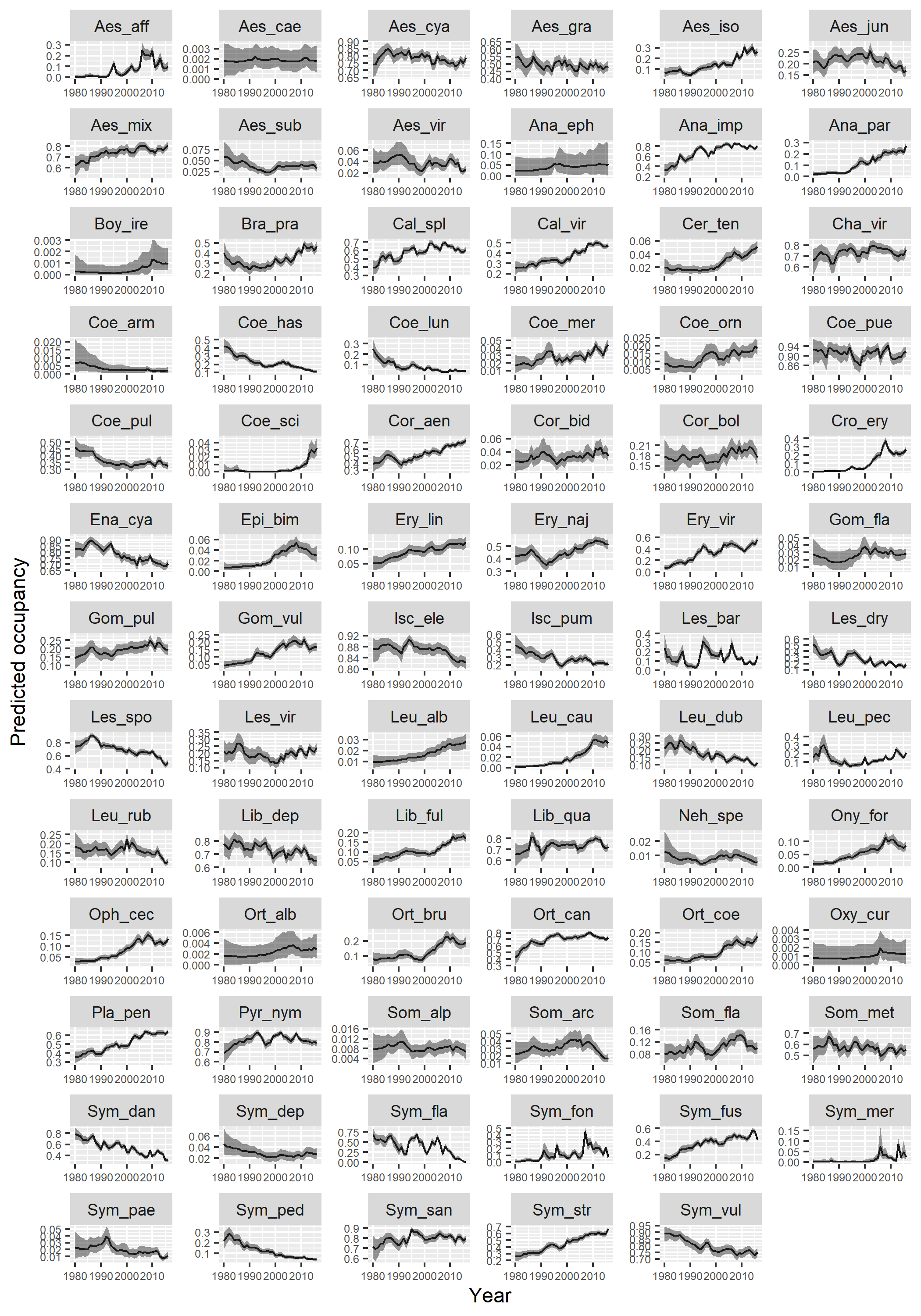

Table S2: Species in each cluster

| Cluster |  |
| --- | --- |
| 1 | Aeshna affinis Aeshna caerulea  Aeshna isoceles Aeshna mixta  Anax ephippiger Anax imperator  Anax parthenope Calopteryx splendens  Calopteryx virgo Chalcolestes viridis  Coenagrion mercuriale Coenagrion ornatum  Crocothemis erythraea Epitheca bimaculata  Erythromma lindenii Erythromma viridulum  Gomphus flavipes Gomphus pulchellus  Gomphus vulgatissimus Leucorrhinia albifrons  Leucorrhinia caudalis Libellula fulva  Libellula quadrimaculata Onychogomphus forcipatus  Ophiogomphus cecilia Orthetrum albistylum  Orthetrum cancellatum Orthetrum coerulescens  Platycnemis pennipes Somatochlora flavomaculata  Sympecma fusca Sympetrum fonscolombii  Sympetrum sanguineum Sympetrum striolatum |
| 2 | Boyeria irene Brachytron pratense  Ceriagrion tenellum Coenagrion scitulum  Cordulegaster bidentata Cordulegaster boltonii  Cordulia aenea Erythromma najas  Orthetrum brunneum Oxygastra curtisii  Sympetrum meridionale |
| 3 | Aeshna juncea Lestes barbarus  Leucorrhinia rubicunda Pyrrhosoma nymphula  Somatochlora arctica |
| 4 | Aeshna subarctica Coenagrion puella  Lestes virens Leucorrhinia pectoralis  Nehalennia speciosa Somatochlora alpestris  Sympetrum depressiusculum |
| 5 | Aeshna cyanea Aeshna grandis  Aeshna viridis Coenagrion armatum  Coenagrion hastulatum Coenagrion lunulatum  Coenagrion pulchellum Enallagma cyathigerum  Ischnura elegans Ischnura pumilio  Lestes dryas Lestes sponsa  Leucorrhinia dubia Libellula depressa  Somatochlora metallica Sympecma paedisca  Sympetrum danae Sympetrum flaveolum  Sympetrum pedemontanum Sympetrum vulgatum |

Fig. S2 - Repeat calculation of the cluster means removing the species with the largest changes in each cluster.

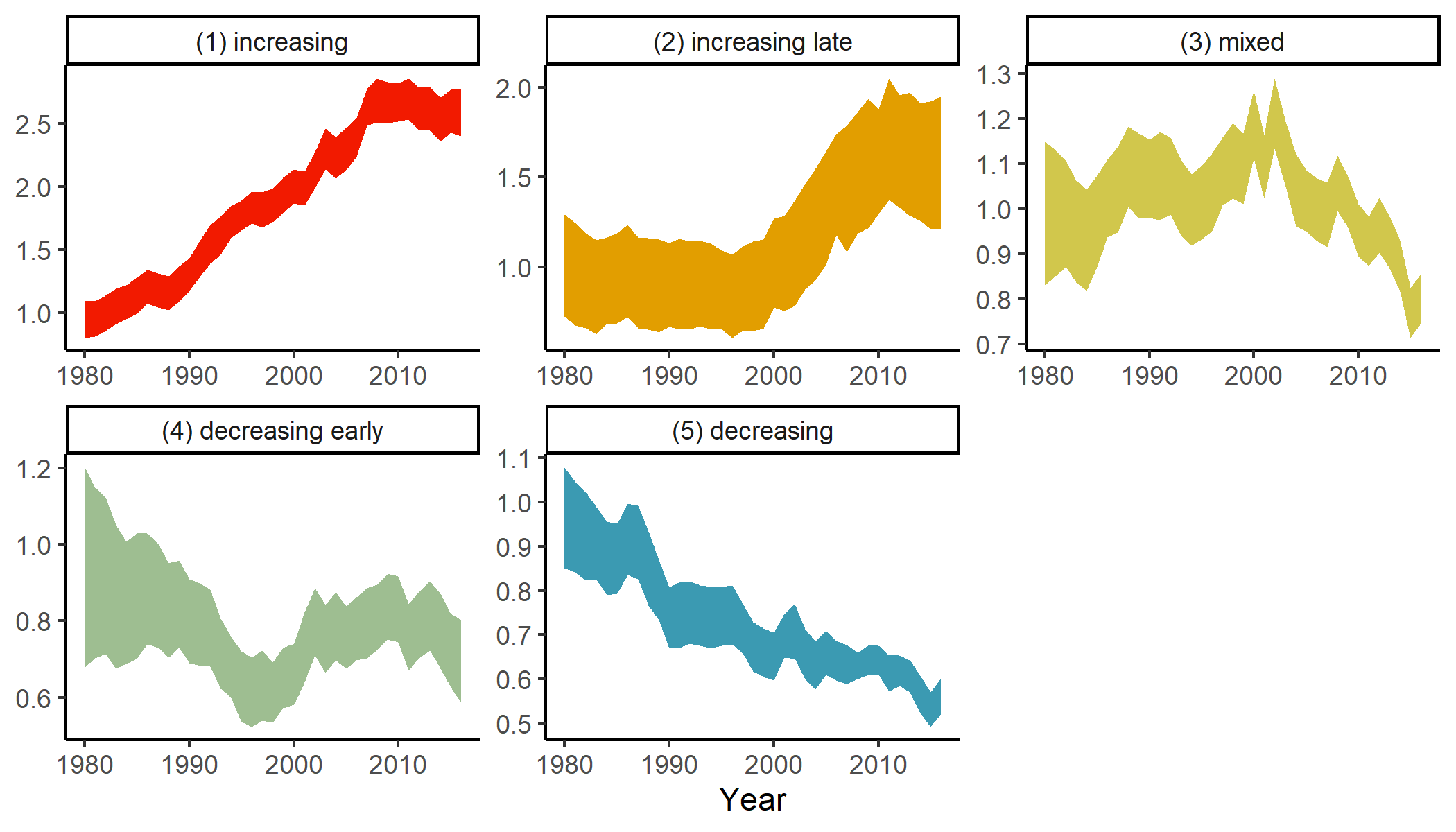
